# Intranasal therapies for neonatal hypoxic-ischemic encephalopathy: Systematic review, synthesis, and implications for global accessibility to care

**DOI:** 10.1101/2024.09.26.615156

**Authors:** Andrew S. Cavanagh, Nazli Kuter, Benjamin I. Sollinger, Khyzer Aziz, Victoria Turnbill, Lee J. Martin, Frances J. Northington

## Abstract

Neonatal hypoxic-ischemic encephalopathy (HIE) is the leading cause of neurodevelopmental morbidity in term infants worldwide. Incidence of HIE is highest in low and middle-income communities with minimal access to neonatal intensive care and an underdeveloped infrastructure for advanced neurologic interventions. Moreover, therapeutic hypothermia, standard of care for HIE in high resourced settings, is shown to be ineffective in low and middle-income communities. With their low cost, ease of administration, and capacity to potently target the central nervous system, intranasal therapies pose a unique opportunity to be a more globally accessible treatment for neonatal HIE. Intranasal experimental therapeutics have been studied in both rodent and piglet models, but no intranasal therapeutics for neonatal HIE have undergone human clinical trials. Additional research must be done to expand the array of treatments available for use as intranasal therapies for neonatal HIE thus improving the neurologic outcomes of infants worldwide.

## 1. Introduction

Neonatal hypoxic-ischemic (HI) encephalopathy (HIE) results from perinatal cerebral oxygen and glucose deprivation. It causes permanent brain injury and is the leading cause of neurodevelopmental morbidity in term infants worldwide^1^. While the incidence of HIE may be as low as 1.5/1000 live births in high-income countries, as of 2010, HIE incidence was as high as 10.4/1000 live births in South Asia and 14.9/1000 live births in Sub-Saharan Africa^2^. It is estimated that South-Southeast Asia and Sub-Saharan Africa contribute to a combined 85% of HIE case incidence worldwide^2^.

In most low and middle-income countries (LMIC), less than 70% of infants are born within a designated medical facility, and less than 5% of these infants have access to a specialized neonatal intensive care unit (NICU)^2^. Although the current standard of care for HIE, therapeutic hypothermia (TH), significantly decreases long-term morbidity and mortality in high-income countries^3^, it has been shown to increase case mortality in LMICs^4^. Due to the high labor and capital investment needed for therapeutic hypothermia, these disparities in mortality may be attributable to both a lack of supportive care and a scarcity of effective cooling equipment^4^.

A lack of access to specialized neonatal care is not confined to LMICs. Even in countries with a more advanced medical infrastructure, many infants continue to live out of reach from centers that can provide acute neurologic intervention^5^. As of 2013, only 50% of micropolitan women and less than 30% of rural women in the United States lived within a 30-mile drive of a NICU^5^. Accessibility to care for infants born in these communities must be considered in the search for novel therapeutic interventions for HIE.

Intranasal (IN) administration of therapies involves delivery of a therapeutic directly into the naris^6^. Recently, IN therapeutic administration has seriously emerged as a viable neuropharmacologic route because of its minimally invasive nature and high nervous system potency^6^. IN administration has especially grown in the delivery of analgesics in many NICUs across the United States^7^. Clinical studies have shown IN sedation to be both safe and effective in term and preterm infants validating the ability to target the central nervous system (CNS) through the neonatal nasal mucosa^7–9^.

Because of the potential of high efficacy and increased accessibility of IN drug delivery for neonates worldwide, we propose that IN treatments may play an important role in expanding the global accessibility of HIE treatment^10, 11^ (Fig. 1). In this review, the nasal anatomy, physiology, and histology relevant to the IN administration of therapies for HIE will be discussed in the context of both human and animal models. The current state of research concerning IN therapies for HIE will be reviewed, and the relative development of experimental IN therapeutics toward large animal models and human clinical trial will be semi-quantitatively evaluated. Finally, the implications of accelerating research on IN treatments towards finding globally accessible novel therapies for HIE will be discussed.

**Figure 1:**
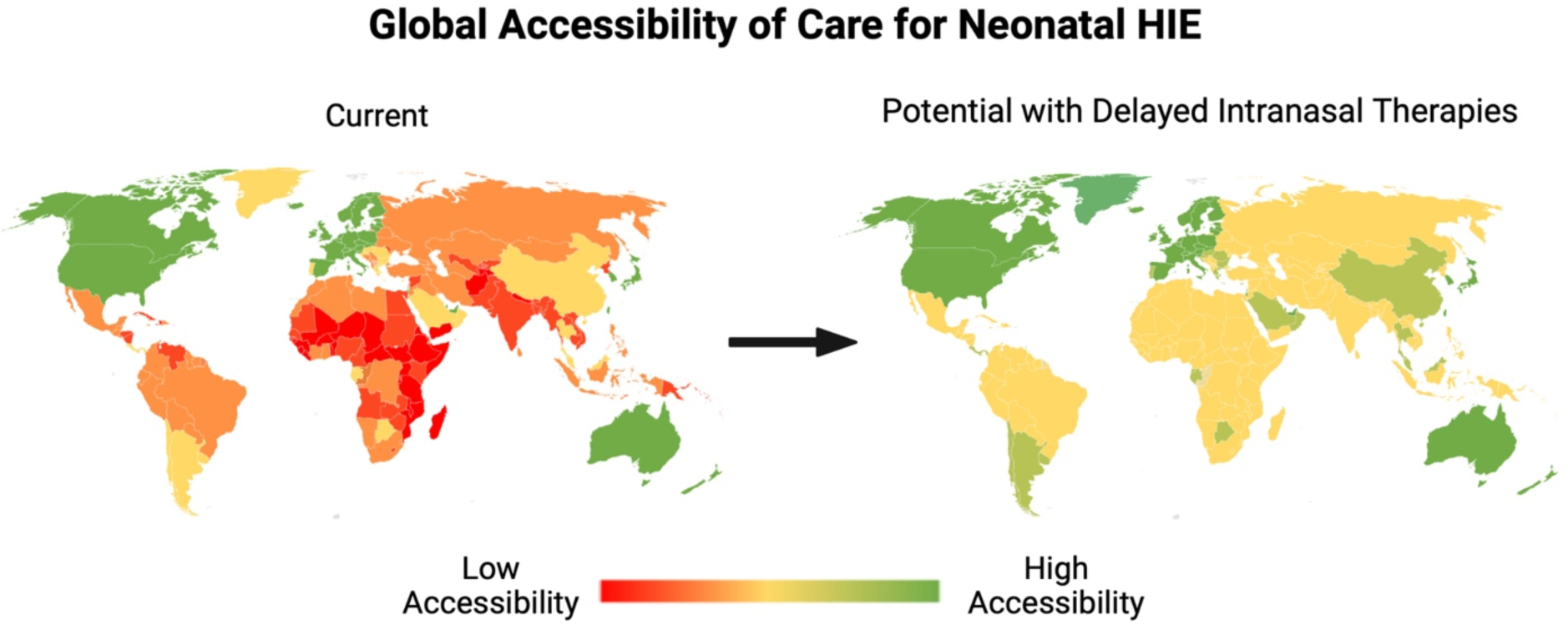
Potential Improvement in Global Accessibility to HIE Care with IN Therapies. Countries with low accessibility to HIE care are shown in red, while countries with accessible HIE care are shown in green. Countries with varying intermediate degress of HIE care accessibility are in yellow and orange. Improvement in accessibility to HIE care is a global issue. Figure adapted from data provided by the World Bank ^96^. Created in BioRender.

### 1.1. Human Nasal Anatomy, Physiology, and Histology

The main functions of the nasal cavity include the passage of air for breathing and the delivery of odorants for olfaction^12^. Its four principal regions, distinguished by their anatomic and physiologic differences, include the nasal vestibule, the nasal atrium, the respiratory region (conchae), and the olfactory area^12, 13^ (Fig. 2). The nasal vestibule is the most anterior region of the nasal cavity^12–14^. The stratified squamous, keratinized epithelium of the vestibule projects hairs, vibrissae, that filter inhaled particles affording rigorous protection to the nasal cavity but foreclosing the delivery of many large-molecule therapeutics^13, 14^. The nasal atrium, the intermediate anatomic region of the nasal cavity, is lined with stratified squamous epithelium anteriorly, while microvillous pseudostratified columnar cells line the posterior section^12, 13^.

**Figure 2:**
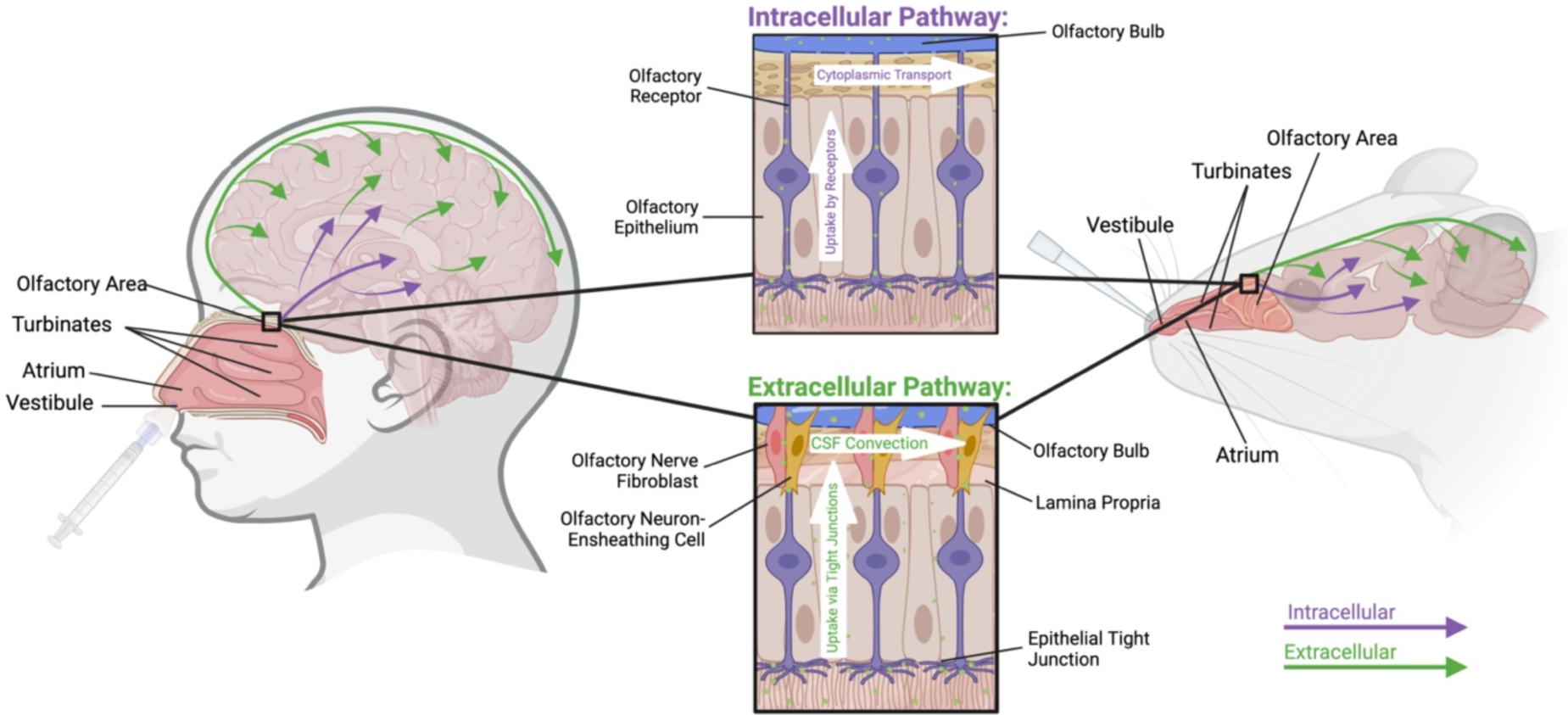
Human and Mouse Nasal Anatomy and Histology. Shown are anatomic and cellular structures of the human and mouse nasal cavity relevant to the intranasal delivery of therapeutics to the CNS ^12, 97^. Both intracellular and extracellular therapeutic pathways are portrayed ^22, 27, 98^. Created in BioRender.

The respiratory region is the largest of the nasal cavity and is divided by superior, middle, and inferior turbinates projecting from the lateral wall responsible for the humidification and temperature regulation of inhaled air^12, 13, 15^. Spaces, meatus, between turbinates bring inhaled air in close contact with the respiratory mucosal surface. The respiratory mucosal surface consists of pseudostratified columnar epithelial cells with their basement membrane and the lamina propria, a network of capillaries, vessels, nerves, glands, and immune cells. The lamina propria may provide the opportunity for IN therapeutics to reach systemic circulation^12, 13^. IN therapeutics also target the olfactory region which lies on the roof of the nasal cavity^12^. Olfactory neuroepithelium provides direct access to the CNS. Pseudostratified epithelium and olfactory receptors projecting through the apical surface afford inhaled substances direct transfer into the CNS^12, 16^.

### 1.2. Mechanism of IN Therapeutic Delivery to the Central Nervous System

The blood-brain barrier (BBB) makes delivery of exogenous substances to the CNS especially difficult^17^. Even for therapeutics capable of permeating the BBB, multidrug efflux protein transporters including P-glycoprotein (P-gp) limit therapeutic permeability^18^. Although olfactory epithelium presents an opportunity to bypass the BBB through direct delivery to the CNS, efflux pumps are still heavily present in olfactory neurons and supporting cells^19^.

Approximately 50% of all neurotherapeutic drug candidates are substrates to P-gp thus narrowing the number of potential IN therapeutics^20^. Nonetheless, IN administration presents a potentially rapid and potent route of CNS exposure for permeable therapeutics in comparison to other practicable routes of administration^18^.

IN therapeutics are delivered to the CNS via intracellular and extracellular routes^17^ (Fig. 2). Drug delivery via the intracellular pathway begins with both endo- and pinocytosis by an olfactory receptor neuron^21^. The therapeutic is subsequently engulfed by an endosome, delivered to the Golgi, and trafficked caudally via axoplasmic transport in both the olfactory and trigeminal nerves. Down the length of the nerve, it is exocytosed into extracellular space and, if engulfed by adjacent cells and further distributed throughout the CNS^21, 22^. Although effective, this slow axoplasmic transport is unlikely to be the major route of drug delivery to the CNS^17, 23^

Extracellular distribution to the CNS is initiated when IN therapeutics cross epithelial tight junctions into paracellular clefts^24^. In the lamina propria, IN therapeutics diffuse through the perineural space between olfactory neuron-ensheathing cells and olfactory nerve fibroblasts allowing access to the meninges and to the subarachnoid space^24–26^. The therapeutic can then be delivered to CNS tissue via CSF convection. Systemic transport via blood and lymphatic vasculature through the lamina propria may also deliver IN therapeutics to the CNS^26,27^.

### 1.3. Considerations for Human Neonatal Nasal Anatomy, Physiology, and Histology

The contributions of neonatal-adult differences in nasal anatomy and physiology towards comparative pharmacokinetic differences of IN therapeutics are relatively unknown. Primary olfactory receptors in the nasal cavity are present by the eighth week of gestation and are fully functional by the twenty-fourth week^28^. Neonates are obligate nose breathers and possess more narrow nasal passages than adults creating a greater baseline nasal air resistance that must be overcome^29^. Since neonates have both an increased nasal resistance and a heightened respiratory rate compared to adults^30^, it is possible that these factors may contribute to an increased absorbance of IN therapeutics^31^. However, since neonates and preterm infants are more susceptible to nasal obstruction than older infants, access of therapeutics to the CNS via the lamina propria and olfactory epithelium may be limited^32^. The lack of knowledge of IN pharmacokinetics in neonates requires that all research be accompanied by direct or indirect proof of CNS delivery of the therapeutic. It is furthermore important to determine if olfactory neuroepithelium is damaged by HIE and if IN drug delivery is subsequently perturbed^33, 34^.

### 1.4. Comparative Nasal Anatomy, Physiology, and Histology of Animal Models

The internal gross anatomy of the nasal cavity of adult rodents and piglets is comparable to that of adult humans^35, 36^. All contain two physiologically symmetric nares that extend caudally towards a nasopharynx^35^. The size of various structures within the anterior nasal cavity differs across species, however, these differences in anatomy likely do not preclude translation of IN pharmacology research across species^34^. More consequential differences lie in the proportional size and innervation density of olfactory areas and the lamina propria^34–36^. Rodents and piglets exhibit a proportionally larger olfactory area and lamina propria than those of humans suggesting that IN therapeutics may reach the brain more potently in rodent and piglet models than in humans^35^.

The intracellular and extracellular routes of drug delivery in animal models are similar to humans^16, 18, 21^. Little research exists, however, on the comparative nasal anatomy of early-life rodents and piglets. Thus, therapeutic trials in rodents and piglets are largely proof of concept that will require additional testing before implementation of IN treatment strategies in humans^34–36^. Fortunately, such testing is already underway^37^.

### 1.5. Optimizing the Accessibility of Therapeutics for HIE

Despite the concentration of HIE burden in LMICs, many issues, both medical and sociopolitical, impede the widespread implementation of various therapies for HIE. For any therapy, there are a variety of factors intrinsic to the therapeutic that heavily impact its accessibility^38^. These include cost of the therapy itself, cost of distribution and refrigeration requirements, its ease of administration, and how quickly, how often, and for how long must it be given^38^. IN therapies would be much more easily administered and repeatedly dosed, if necessary, than other currently available treatments for HIE^6, 18, 39, 40^. Given IN treatment does not require intravenous (IV) access, it poses less risk for adverse complications related to IV administration^41^.

While effective therapeutic hypothermia must be initiated within the first 6 hours of birth^42^, many communities lack the prehospital transportation infrastructure to initiate any intervention in that timeframe^43^. Alternatively, many potential IN therapeutics target later phases of injury and offer a regenerative approach that could afford administration days after injury onset^37, 44–51^.

### 1.6. Safety Concerns for the IN Administration of Therapeutics to Neonates

Physical and chemical irritation of nasal epithelium is a major concern with the IN administration of therapeutics^34^. Nonetheless, irritation can often be avoided through careful and precise delivery of the therapeutic to the nasal cavity^52^. Coadministration of lidocaine has been reported for this purpose, however, this should be considered carefully for neonates^53^. Drug dose determination is also of concern for the IN administration of therapeutics as recommended systemic doses may need to be altered greatly across routes^54, 55^.

The IN administration of analgesics and sedatives are well studied in human neonates suggesting IN delivery to neonates is safe and effective^7–9^. In a systematic review including 400 neonates, IN administration of analgesics and sedatives was associated with a significant reduction in markers of pain, measured by clinical scales, skin conductance, heart rate, and crying time, compared to other routes of sedation^7^.

## 2. State of the Art Intranasal Therapies for HIE

The idea of using an IN route to treat HIE is not new. In fact, the first report of IN administration of a therapeutic for HIE was published in 2003 on the delivery of Insulin-like Growth Factor 1 (IGF-1) in rodents^56^. We compiled relevant literature via an internet search of the phrases “Neonatal Hypoxic-Ischemic Encephalopathy Intranasal” and “Neonatal Brain Intranasal” on Google Scholar and PubMed search engines. All literature appearing within the first 10 original search results, sorted by relevance, for each criterion that pertained to the IN delivery of an experimental therapeutic for HIE was included (Table 1). In addition, the human clinical trial concerning perinatal arterial ischemic stroke was included despite it not being a study of HIE^37^.

**Table 1:**
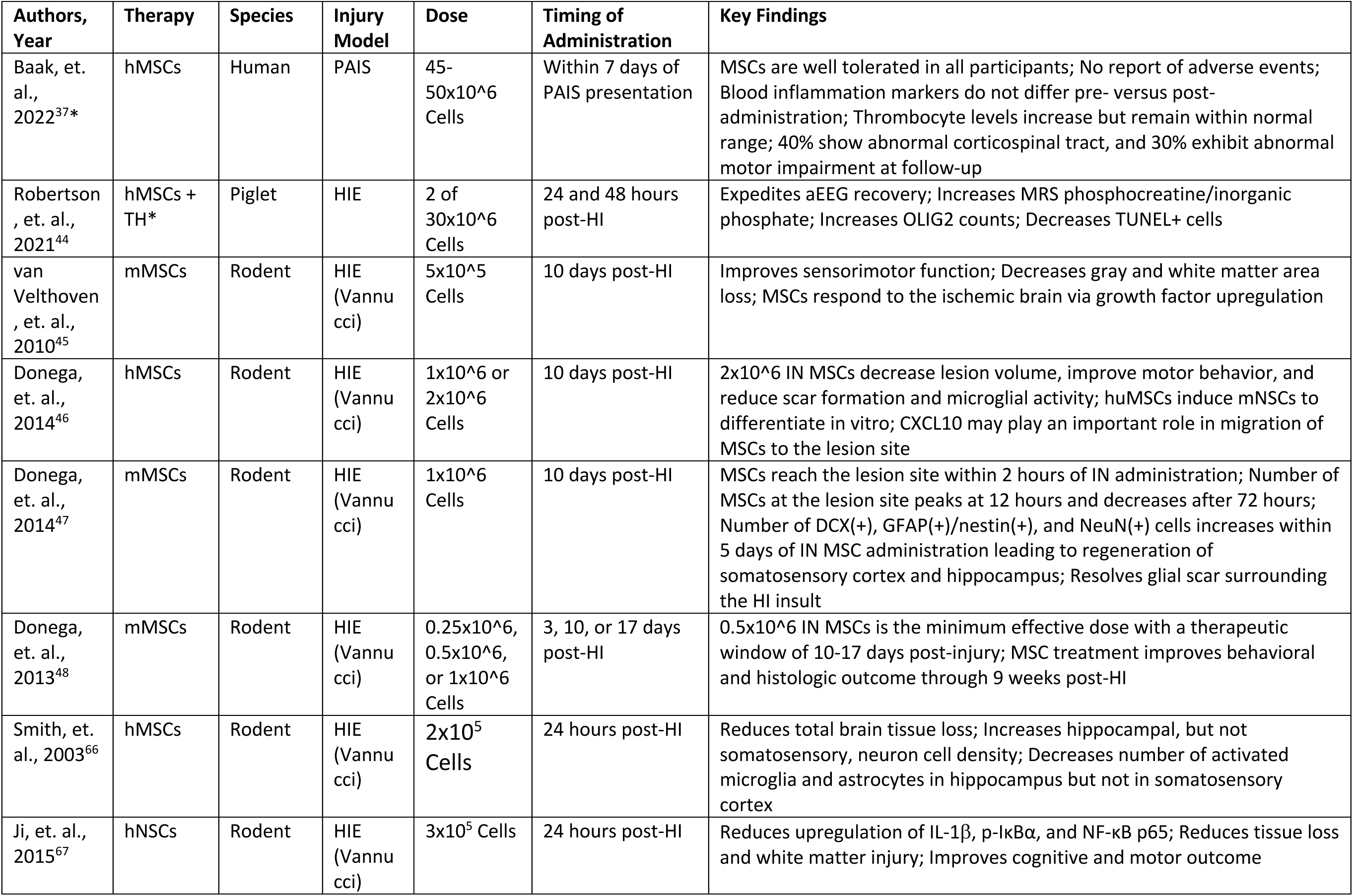
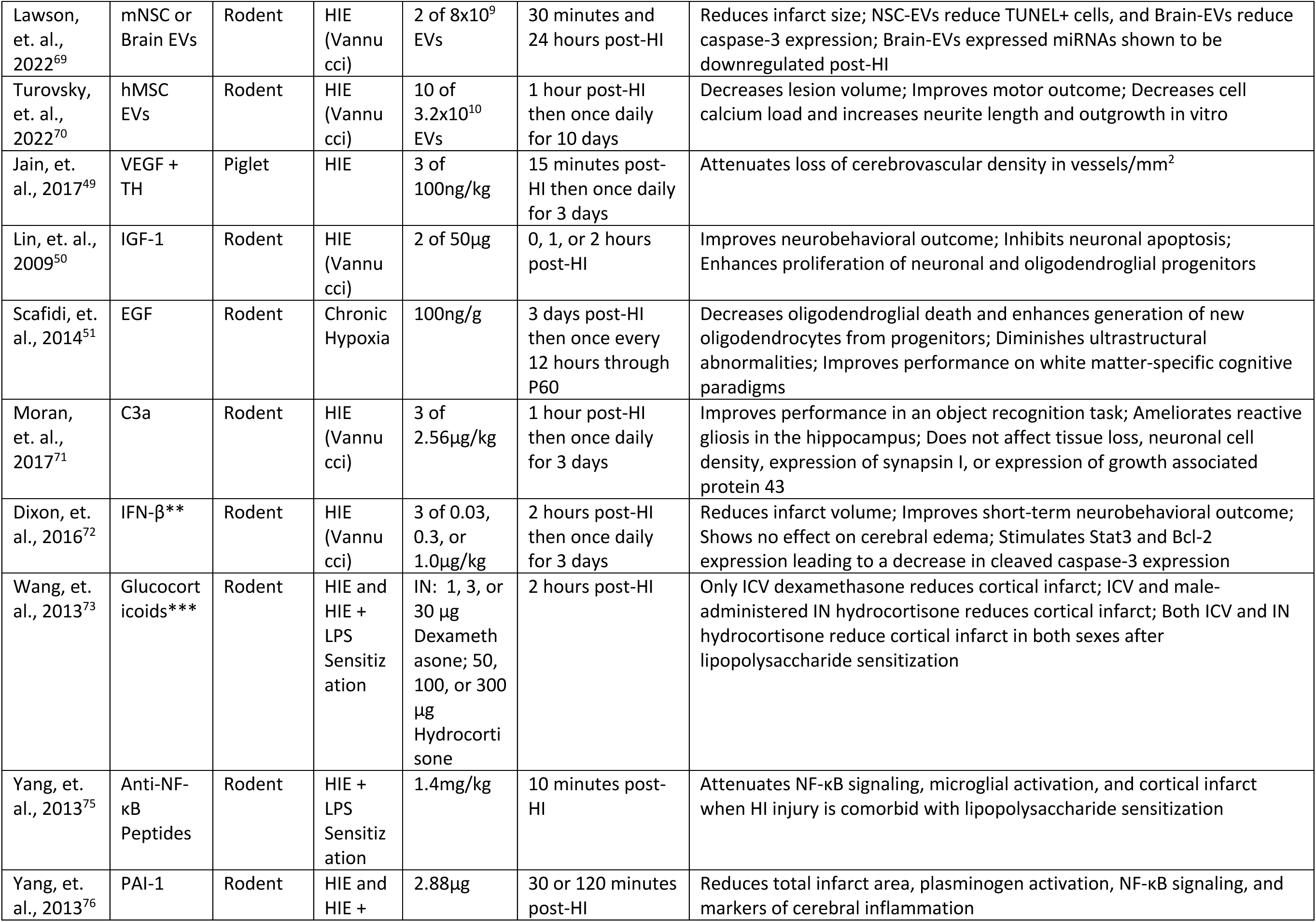
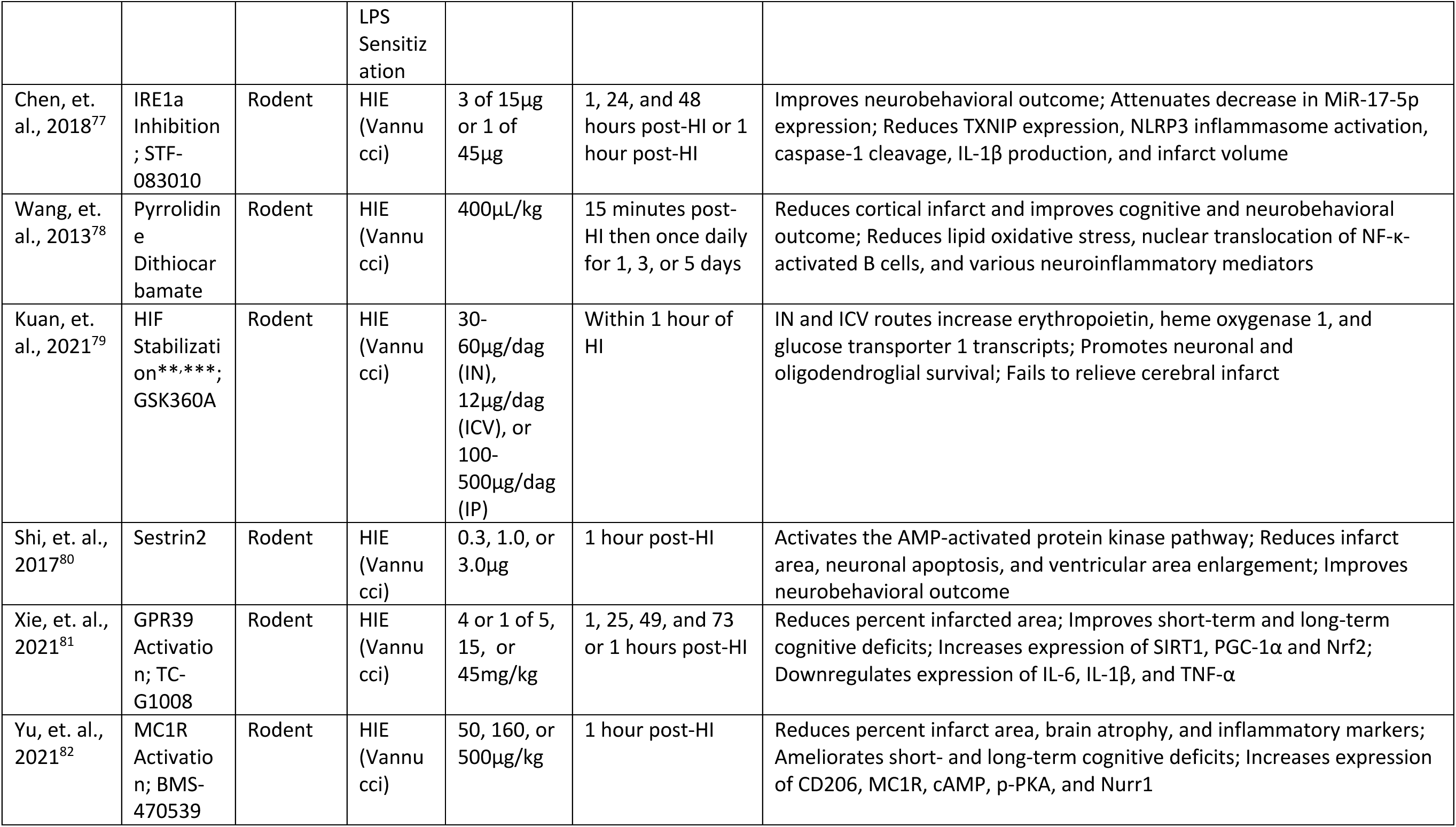
Literature Reviewed. In addition to IN administration: *Study also assessed IV administration of the therapeutic, **Study also assessed IP administration of the therapeutic, ***Study also assessed ICV injection of the therapeutic

### 2.1. IN Administration of Therapeutics for Human Neonatal Brain Injury

Although no IN therapeutics have been translated to human clinical trial for HIE, IN therapeutics have been evaluated in human neonates for ischemic brain injuries including stroke^37^.

### Mesenchymal Stromal Cells for Perinatal Arterial Ischemic Stroke

An early-stage clinical trial is complete evaluating the safety and feasibility of IN mesenchymal stromal cells (MSC) for perinatal arterial ischemic stroke (PAIS)^37^. Ten full-term (≥36 week gestation) neonates with MRI-confirmed PAIS in the middle-cerebral artery region and pre-Wallerian changes in the corticospinal tract received 45-50x10^6^ bone marrow-derived MSCs IN within 7 days of stroke presentation. In this phase I study, IN administration of MSCs was well tolerated in all ten neonates with no adverse events requiring clinical intervention reported. Platelets increased, but not outside of normal range, and there were no changes in blood inflammation markers (C-reactive protein, procalcitonin, and leukocyte count) upon administration of MSCs. Follow-up MRI at 3 months did not show any unexpected cerebral abnormality, and only four patients showed an asymmetric corticospinal tract on 4-month follow-up MRI. Abnormal motor assessments were found in three infants at 4 months^37^.

Despite this being a trial in neonates with PAIS, rather than HIE, it supports the safe IN administration of MSCs for human neonatal brain injury in an acute care setting.

### 2.2. Administration of Therapeutics in Piglet Models of HIE

Swine models are excellent models for HIE^57, 58^. Phylogenetically, the pig is estimated to be threefold closer to humans than mice^57^. Moreover, the gyrencephalic pig brain is more similar in anatomy and development to humans than is the rodent brain^59^. For these reasons, swine models are thought to be ideal for drug discovery^57^.

#### Mesenchymal Stromal Cells

The efficacy of human (h) MSCs adjunct to therapeutic hypothermia for HIE in a swine model was tested with either IV or IN MSCs both in two doses of 30x10^6^ cells 24- and 48-hours post-HI^44^. Amplitude-integrated electroencephalogram (aEEG) recovery was more rapid and phosphocreatine/inorganic phosphate ratio on MRS was higher after 48 hours when animals were administered IN MSCs with TH compared to HI or HI with TH, respectively. OLIG2 counts of oligodendrocytes in the hippocampus, internal capsule, and periventricular white matter were higher in the IN MSC group than the IV MSC group. TUNEL-positive cells were reduced in the internal capsule in the IN MSC with TH treated group compared to TH alone. IN MSC delivery efficacy was tested with PKH26-labeled MSCs which were detected in the brain 12 hours after administration^44^.

#### Vascular Endothelial Growth Factor

Rh-VEGF165 was administered IN to piglets at a dose of 100ng/kg 15 minutes after HI then daily for three days^49^. IN Rh-VEGF165 resulted in recovery of cerebrovascular density (measured in vessels/mm^2^) from that of injured controls and comparability to shams without promoting neovascularization^49^.

### 2.3. IN Administration of Therapeutics in Rodent Models of HIE

Rodents provide an efficient mode to study basic mechanistic, biochemical, histologic, and behavioral outcomes of HIE^60^. Regarding the IN administration of therapeutics, there are similar routes of intra and extra-cellular transport from the nares to the brain in rodents and humans, allowing for determination of whether a therapy is brain penetrant when given intranasally^17^. Rodent models are otherwise well studied, widely used, afford access to numerous genetically modified models, have shown partial neuroprotection from TH, and afford opportunity for behavioral testing with numerous rigorous, high throughput protocols^60–64^. Unless otherwise noted, the HI injury model used in the following literature was the Vanucci model of neonatal HIE^65^.

#### 2.3.A. Cell-Based Therapies Mesenchymal Stromal Cells

van Velthoven, et. al.^45^ pioneered use of IN MSCs for neonatal HI by delivering 5x10^5^ PKH-26-labeled mouse bone marrow-derived MSCs IN. Undifferentiated MSCs remained detectable 28 days after injury and treatment in the ipsilateral hippocampus. IN MSC treatment improved sensorimotor function 21 days and 28 days post-HI and decreased gray and white matter area loss by 34 and 37%, respectively. MSC administration altered the toxic milieu in the ischemic region through upregulation of fibroblast growth factor 2 and nerve growth factor^45^.

To further evaluate the translational relevance of cell-based therapies as a regenerative therapeutic strategy for HIE, Donega, et. al.^46^ evaluated the IN administration of hMSCs with 1x10^6^ or 2x10^6^ cells to postnatal day (P) 9 rats after P9 HI injury. The larger dose of hMSCs decreased lesion volume, improved motor behavior, and reduced scar formation and microglial activity. The chemotactic factor CXCL10, released by both neurons and glia after HI injury, attracted MSCs to the lesion site. Furthermore, hMSCs induced differentiation of mouse (m) neural stem cells (NSC) in an in vitro differentiation assay^46^.

Using MRI and immunohistochemistry to explore the pathobiology and pharmacokinetics of IN mMSCs, Donega, et. al. showed that MSCs reached the lesion site as early as 2 hours after administration and that the number of MSCs peaked at 12 hours after administration and decreased sharply after 72 hours^47^. At the lesion site, GFAP^+^/nestin^+^ and DCX^+^ expression increased from 3 to 5 days after administration with an increase in number of NeuN^+^ neurons within 5 days. Dramatic regeneration of the ipsilateral somatosensory cortex and hippocampus occured by 18 days after MSC administration. There was resolution of the glial scar surrounding the injury measured with a decrease in reactive astrocytes and an increase in the M2 microglia phenotype^47^.

Study of the long-term efficacy of IN mMSCs found that a single dose of 0.5x10^6^ MSCs was the minimally effective dose to decrease infarct size and improve social discriminatory and sensorimotor behavioral outcomes beginning 10-17 days post-injury and maintained for 9 weeks post-injury^48^.

Moreover, 2x10^5^ human umbilical cord derived MSCs delivered IN 24 hours post-HI reduced total brain tissue loss, while hippocampal, but not somatosensory, cortical neuron cell density increased ^66^. Confirming the neuropathology results of Donega et. al.^47^, MSCs decreased the number of activated microglia and astrocytes in the hippocampus but not in the somatosensory cortex^66^.

#### Neural Stem Cells

3x10^5^ PKH-26-labeled hNSCs administered 24 hours after P7 HI injury in rats were detectable in the brain 24 hours post-administration and reduced upregulation of IL-1β, p-IκBα, and NF-κB p65^67^. Tissue loss and white matter injury were reduced 42 days post-HI, and there was improved performance on righting reflex, gait, grid, social choice, and water maze tests^67^.

Interestingly, hypoxic pretreatment of intranasal CM-Dil-labeled hNSCs prior to delivery improved delivery efficacy of the hNSC to the brain^68^. Mechanisms may include hypoxia-enhanced expression of CXC chemokine receptor 4 and improved migration of NSCs to the lesion. Additionally, delivery with a an intranasal catheter, as opposed to inhalation, and repeated dosing at intervals ≥ 24 hours also improved delivery of cells^68^.

#### Extracellular Vesicles

To evaluate the efficacy of IN extracellular vesicles (EV) for neonatal HI, 8x10^9^ NSC EVs or hypoxia-preconditioned brain EVs were administered IN 30 min and 24 hours post-HI. Infarct size was reduced following administration of either EV^69^. NSC-EVs reduced expression of TUNEL+ cells, however only brain-EVs reduced caspase-3 expression. The hypoxia preconditioned brain-EVs also contained miRNAs known to be downregulated after neonatal HI, miR-342-3p and miR-330-3p^69^. In a further study, administration of 3.2x10^10^ MSC-derived EVs given 1 hour following P7 HI and then daily for 10 days improved lesion volume at 60 days post injury^70^. Lesion volumes of IN MSC-EV-treated rats were no different than non-treated HI rats at 40 days post injury. IN MSC-EVs ameliorated sensorimotor dysfunction in fore- and hindlimbs on day 40 and 60 post-HI. In vitro, MSC-EVs decreased the hypoxic-ischemic calcium load of neuroglia and increase neurite length and outgrowth. MSC-EV-induced inositol triphosphate receptor-related calcium oscillations were associated with resistance to calcium overload in both neurons and glia, suggesting that PI3K signaling is necessary for MSC-EV protection of HI injured neural cells^70^.

### 2.3. B. Growth Factor Regulation Insulin-Like Growth Factor 1

At 0, 1, or 2 hours following HI injury, 2 50µg doses, 1 hour apart, of human recombinant (rh) IGF-1 administered IN resulted in a high concentration of rhIGF-1 detectable in the brain 30 minutes following administration^50^. Cortical rhIGF-1 reduced apoptotic death via phosphorylation in the Akt signal transduction pathway and enhanced proliferation of neuronal and oligodendroglial progenitors. Performance on a passive avoidance task improved with IN rhIGF-1 after HI^50^.

#### Epidermal Growth Factor Receptor Signaling

Scafidi, et. al.^51^ evaluated the efficacy of IN heparin binding epidermal growth factor (EGF) to enhance the endogenous response of EGF receptor-expressing progenitor cells. IN treatment with 100 nanograms/gram IN heparin-binding EGF every 12 hours through P60 for repeated hypoxia from P3 through P11 resulted in heparin-binding EGF and phosphorylated EGF receptor detectable in white matter from 5 to 30 minutes following IN administration. IN heparin binding EGF decreased oligodendroglial cell death and enhanced generation of new oligodendrocytes from progenitor cells. IN heparin binding EGF decreased ultrastructural abnormalities in corpus callosum, cingulum, and external capsule, and improved motor coordination ^51^. This study of repetitive hypoxic injury in a preterm model demonstrates the feasibility of IN delivery of both late and chronic therapies for hypoxia-related brain injuries.

### 2.3. C. Immune System Modulation C3a

After P9 HI injury, mouse pups were administered 2.56µg/kg C3a IN to determine if modulation of the complement cascade is neuroprotective^71^. IN administration of C3a started one hour after HI and continued daily for three days. C3a treated mice exhibited improved long-term memory post-HI compared to vehicle-treated controls. IN C3a ameliorated HI-induced reactive gliosis in the hippocampus, but did not improve hippocampal tissue loss, decreases in neuronal cell density, and expression of synapsin 1 or growth associated protein 43^71^.

#### Interferon Beta

In a comprehensive set of experiments, Dixon, et. al.^72^ explored dose optimization, pharmacokinetics, neuropathology, and early neurobehavior following IN administration of IFN-β for neonatal HIE. 3 doses of 0.03, 0.3, or 1.0 µg/kg of recombinant human IFN-β were administered IN beginning two hours after P10 HI injury and continuing daily for three days.

Cortical IFN-β exceeded that of vehicle-treated controls by 1 hour, peaked at 12 hours, and began to fall at 24 hours post-administration. The 2 highest doses of IN IFN-β reduced total brain infarct volume in HI-injured rats. Due to increases in P-STAT3 and Bcl-2 expression and decreases in cleaved caspase-3 expression, the authors surmised that IN IFN-β treatment blocked injury-induced apoptosis. IN IFN-β did not ameliorate development of consistent, rapid evolution of edema as injury is initiated. IN administration of IFN-β related to an improvement of performance on surface righting and negative geotaxis reflex tests up to 72 hours post-injury^72^.

#### Glucocorticoids

Two drugs (hydrocortisone and dexamethasone), two dosing routes (IN and intracerebro-ventricular (ICV), and 3 IN doses (1, 3, or 30µg dexamethasone; 50, 100, or 300µg hydrocortisone) and 1 ICV dose (0.1µg dexamethasone; 10µg hydrocortisone) were used for treatment of HI or HI with lipopolysaccharide sensitization (LPS)^73^. In the HI model, both drugs given ICV decreased mean infarction size, while only males received benefit from the highest tested dose of IN hydrocortisone. In contrast, both drugs given via both routes were therapeutic in the setting of HI and LPS sensitization^73^

#### Anti-NF-κB Peptides

In the LPS sensitized HI model, IN administration of Tat-NBD, a peptide-inhibitor of Nuclear Factor- κB (NF-κB) activation and nuclear translocation, was given 10 minutes after P7 HI injury at a substantially lower dose (1.4 mg/kg) than is neuroprotective when administered IV^74, 75^. Tat-NBD was detectable by 30 minutes in the olfactory bulb and in the cerebral cortex by 60 minutes. IN Tat-NBD reduced cortical NF-κB activation after LPS sensitized HI injury.

Treatment reduced total infarct volume, cortical microglial activation, and neuronal apoptosis^75^.

### 2.3. D. Protease Inhibitors

#### Plasminogen Activator Inhibitor-1

2.88µg of IN CPAI-1, a stable mutant form of PAI-1, given at either 30 or 120 minutes following P7 HI decreased total infarct area at P14^76^. CPAI-1 levels peaked in the olfactory bulbs at 1 hour and in the cerebral cortex at 2 hours after IN administration. HI injury accelerated IN CPAI-1 transfer from the olfactory bulbs to the cerebral cortex. Confirming specificity of drug action, IN CPAI-1 decreased activity of both tissue and urinary-type plasminogen activators in the ipsilateral hemisphere at 4 hours after HI and blocked the increase in matrix metalloproteinase-9 activity at 24 hours after HI. In the LPS sensitized model, IN CPAI-1 inhibited induced NF-κB activation at 4 hours after injury and blocked increases in IL-6, Ccl-2 and Tspo, a translocator protein indicative of activated microglia, activation at 24 hours^76^.

### 2.3. E. Cellular Stress Signaling IRE1α Inhibition

After HI injury, IN STF-083010, an IRE1α RNase inhibitor, given in either one dose of 45 µg at 1-hour post-injury or 3 doses of 15 µg at 1, 24, and 48 hours post-injury, showed efficacy in reducing total infarct volume^77^. This attenuation was maintained over a 48-hour span with a multiple-dose paradigm as with a single dose. Improved neuropathology was associated with improvement on lateral pressure, proprioceptive limb placement, forelimb placement, lateral limb placement, and T-maze memory tasks. IN STF-083010 reduced expression of TXNIP and NLRP3 leading to a decrease in glial activation. These effects took place through miR-17-5p^77^.

### Pyrrolidine Dithiocarbamate

Treating acutely at 15 minutes after HI then once daily for 3 days with IN PDTC dose-dependently reduced cortical tissue loss with an effective dose (ED)50 at 27mg/kg^78^.

Neuroprotection was maintained when the initial dose is delayed to 45 minutes after HI. IN PDTC attenuated HI-induced lipid oxidative stress and nuclear translocation of NF-κB and reduced expression of IL-1β and TNFα. The antioxidant and anti-inflammatory effects and neuroprotection seen were associated with improved performance on rotarod, Barnes maze, and context- and tone-related fear conditioning tests^78^.

### Hypoxia-Inducible Factor Stabilization

GSK360A, a potent prolyl-4-hydroxylase inhibitor and hypoxia-inducible factor (HIF) stabilizer, administered ICV, intraperitoneal (IP), or IN at a dose of 12µg/decagram (dag), 100- 500µg/dag, or 30 or 60µg/dag by 1 hour after HI triggered upregulation of erythropoietin, heme oxygenase 1, and glucose transporter 1 transcripts via IN and ICV routes^79^. IN administration reduced cortical neuroinflammatory markers (Tspo, Opn, and TNFα mRNAs) and infarct area^79^.

#### Sestrin2

IN recombinant human Sestrin2 at 0.3, 1.0, or 3.0µg given 1 hour post HI upregulated AMPK phosphorylation and inhibited mTOR. The two larger doses reduced infarct area^80^.

Neuroprotection was accompanied by improvement on righting and geotaxis reflex, rotarod, and water maze tests^80^.

### 2.3. F. Anti-Inflammatory GPCR Activation

Repeated IN dosing to mitigate HI injury was attempted with 5, 15, or 45mg/kg 1 hour post-HI or 15 mg/kg 1, 25, 49, and 73 hours post-HI of IN TC-G1008, an activator of G-protein receptor (GPR)39, a receptor with anti-inflammatory effects^81^. Activation of GPR39 increased expressions of SIRT1, PGC-1α, and Nrf2, showing activation of the SIRT1/PGC-1α/Nrf2 pathway, and decreased expressions of IL-6, IL-1β, and TNF-α. Low and medium dosage of IN TC-G-1008 decreased brain infarct area, and medium dosage improved performance on righting reflex, water maze, and rotarod tests^81^.

IN BMS-470539, an agonist of inflammatory mediator melanocortin-1 receptor (MC1R) increased cAMP and Protein Kinase A activation and Nurr1 production while increasing expression of CD206 (a microglial M2 marker) in neonatal rat HI^82^. It also decreased expression of IL-1β, IL-6, and TNF-α while reducing cortical infarct size. 50, 160, or 500µg/kg BMS-470539 was given one hour after HI. An improvement in performance on geotaxis reflex, foot fault, rotarod, and Morris water maze tasks followed 160µg/kg administration^82^.

## 3. Index of Relative Therapeutic Development

To focus and spur development of novel IN therapeutics for HIE, we propose an index of relative therapeutic development for the therapies reviewed herein. Our model, termed the relative development index (RDI) quantifies the efficacy and the developmental progress of IN therapeutics towards demonstrating a positive result on a variety of common experimental outcomes. The RDI may be used as a resource to focus widespread research effort on IN therapeutics that are both effective and well-developed to provide basis for timely study in large animal models and human clinical trials.

A point-based scoring system was devised to compare the relative development of therapeutics reviewed. Experimental outcomes were extracted from the published studies, and each therapeutic was assigned a score for each outcome of 1 if a positive result was demonstrated or 0 if a negative result was demonstrated or if the outcome is not measured.

Each repetitive demonstration of a positive outcome was given a compounding half point value. Experimental outcomes were then grouped into broader classifications. Points were summed within individual outcome classifications and were multiplied by a weighting factor unique to the classification. The authors determined experimental outcome classifications and weighting factors based on the importance of each factors to accelerate further testing of therapies in large animal studies and human clinical trial. The weighting is displayed in Supplementary Table 1 and were as follows in order of assigned importance: evidence for demonstrated brain penetrance with IN dosing, positive neurobehavioral outcome, improved histomorphologic outcome, evidence of reduced inflammation, and evidence for inhibiting or stimulating a specific molecular mechanism known to affect the HI-injured brain. The scores for all outcome classifications across individual therapies were summed to determine a total relative development score.

Results of the RDI are found in Figure 3. MSCs and IGF-1 obtained the two highest index scores with values of 50.5 and 30.0, respectively. The median RDI score was 16.5 with an interquartile range of ± 7.50.

**Figure 3.**
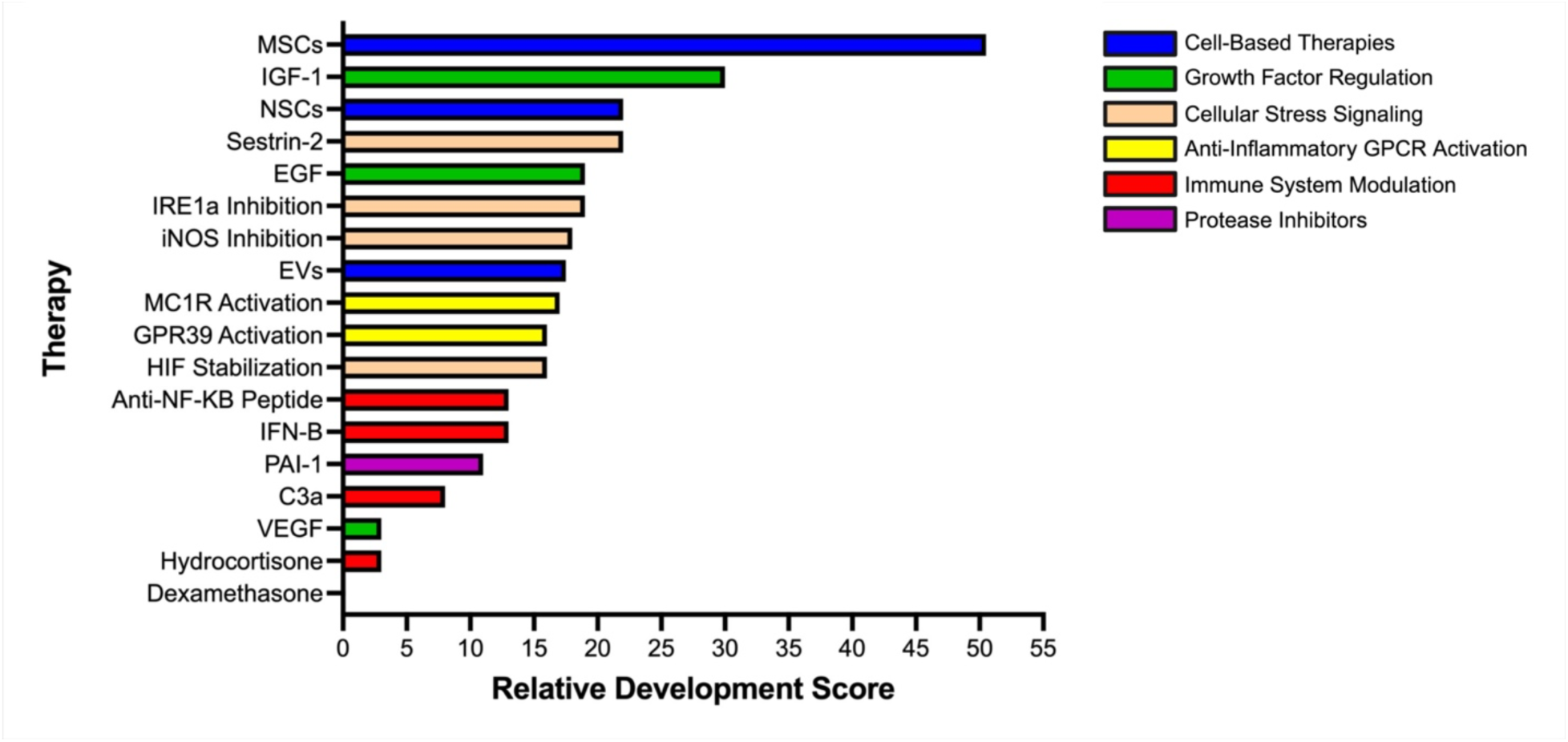
IN Therapeutic RDI. Therapies are indexed according to the method described in Section 3. Therapies similar in mechanism are color-coded according to the legend on the right.

## 4. Discussion

Globally, HIE is the fifth leading cause of death in children under five years of age, however, most patients do not have access to adequate life-saving intervention^2, 83^. Innovative therapeutic options for infants with HIE in both developed and developing communities are urgently needed.

IN therapeutic strategies pose an eminently accessible approach to HIE intervention. They are non-invasive and are easily administered by both skilled and unskilled providers^6, 52^. Many potential IN therapeutics are less expensive than current treatment options and can permeate the CNS potently with a relatively low dose^75, 84^.

The Phase I clinical trial of IN administration of MSCs for PAIS is promising for HIE as it provides precedent for the safe administration of IN therapeutics to the ischemic human neonatal brain^37^. Nonetheless, no human clinical trial of an IN therapeutic for HIE has been done, highlighting the intense need for focused translational investigation.

Laudably, many investigators show evidence of IN drug delivery to the injured brain, mechanistic confirmation of therapeutic action, and one or more biochemical, pathologic, and neurobehavioral improvements in animal models of HIE. Additionally, several show evidence for prolonged neuroprotection and restoration of function following IN treatment. These are ideal preclinical studies from which IN treatments with positive outcomes can be chosen for more advanced studies.

Abundant animal research is ongoing studying the IN administration of cell-based therapies for HIE^44–48, 67, 69^ However, the cognitive and behavioral outcomes of such therapies need to be further delineated. Furthermore, the financial burden and commercial pharmaceutical support compulsory to cell based therapies calls its accessibility to underdeveloped communities into question^85^. These concerns should be addressed before these types of therapies emerge as a widespread clinical intervention for human neonates with HIE.

Related therapies showing promising neuroprotective capacity include a range of growth factor treatments. While VEGF and EGF receptor signaling both demonstrate several positive restorative outcomes, investigation of these and related treatments should include attempts to expand the therapeutic window for each, as the opportunity for intervention in the tertiary phase of HI brain injury is a great need^86^.

Many thematic trends emerge from the broad array of experimental IN therapeutics being tested for neonatal HI. One of the most prominent is the viability of inhibition of NF-κB translocation to the nucleus and prevention of subsequent cytokine production^74, 75, 87^. Given that NF-κB nuclear translocation also evokes pro-survival mechanisms, benefit may be derived from critical consideration of treatment timing. Another prominent trend is regulation of the apoptosome by reduced caspase activation and upstream modulation of substrates such as Bcl- 2 and Protein Kinase B^88^. As chronic HI injury is governed by various forms of programmed cell death, inhibition of apoptosome, necrosome, and inflammasome activity may be crucial strategies for amelioration of late-stage encephalopathy^89–92^.

The RDI attempts to provide a systematic method for investigators to determine where to focus efforts amidst the development of novel IN therapeutics. Cell-based therapies and growth factor regulation demonstrate promising effective development with stem cell therapies and IGF-1 providing the most therapeutic evidence to date. Therapies targeting cellular stress signaling, particularly Sestrin2, also demonstrate highly effective development. Because chronic HI injury may also adhere to neuron-autonomous forms of cell death, therapies targeting growth factor regulation and cellular stress signaling may provide an ideal method to attenuate late-stage injury^89, 91, 93^.

Anti-inflammatory and immunomodulatory targets do not yet demonstrate effective therapeutic development with IN approaches according to the RDI. These studies have focused largely on demonstrating evidence of anti-inflammatory and immunomodulatory outcomes which receive a lower weighting than behavioral and neuropathologic outcomes. Future investigation of these therapies must provide evidence of behavioral deficit amelioration and functional histologic repair to warrant proceeding to larger animal models and to human clinical trials.

Limitations of the RDI, introduced in this review, center primarily on the assignment of weighting factors to outcome classifications, which were ranked by the authors on their perceived importance to moving a therapy forward. Furthermore, the RDI is ignorant to differences in outcome attributable to variation in HI injury model, therapeutic dosing paradigm, or other extraneous factors.

Much of the array of therapies shown effective in the reviewed literature will not translate to effective, stand-alone clinical treatments for neonatal HIE. As is abundantly clear in previous investigation, a single therapy that does not provide a multipronged mitigation of the plethora of injury pathways initiated by neonatal HI injury is unlikely to translate clinically. Thus, a “cocktail” of therapies may be needed to provide optimal treatment^94^. The robust array of treatments under investigation is highly encouraging that an efficacious “cocktail” of IN treatments can be developed.

Research of IN therapeutics has already begun to address a variety of issues that make current treatment options inaccessible to underdeveloped communities. However, there has been limited progress in expanding the proposed therapeutic window of experimental therapeutics after HI injury onset^95^. Many current experimental therapeutics require administration within hours after injury onset. This limits accessibility to communities with delayed access to neonatal neurologic specialization. Expansion of the therapeutic window for HIE as it benefits underdeveloped communities is of utmost importance in future investigation^95^.

In coming years, we expect IN therapeutics for HIE to advance to study in large animal models and begin human clinical trials. With persistent optimization of experimental therapeutics for their global accessibility, IN therapeutics can play a vital role as a safe, effective, and accessible treatments for HIE.

## Acknowledgements

Andrew S. Cavanagh is supported by an ASPIRE award from the Johns Hopkins Krieger School of Arts and Sciences. Frances J. Northington is supported by NS123814, HD110091, HD074593, NS126529, by an Innovations Grant from the Johns Hopkins Department of Pediatrics, and by CDC NU01DD2023000127. Nazli Kuter is supported by a Marshall Klaus Award from the AAP and by a Bauernschmidt Fellowship from the Johns Hopkins School of Medicine Dept. of Pediatrics. Lee J. Martin is supported by NS079348, HD074593, NS123814, and the Johns Hopkins University Alzheimer’s Disease Research Center (AG005146).

---

**Supplementary Table 1.**
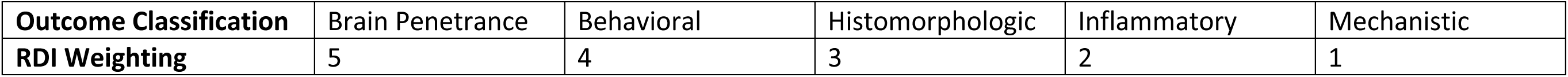
Outcome classifications and respective RDI weightings of the RDI.

## Literature Cited

1. Kurinczuk JJ, White-Koning M and Badawi N. Epidemiology of neonatal encephalopathy and hypoxic-ischaemic encephalopathy. Early Hum Dev 2010; 86: 329–338. 20100616. DOI: 10.1016/j.earlhumdev.2010.05.010.

2. Lee AC, Kozuki N, Blencowe H, et al. Intrapartum-related neonatal encephalopathy incidence and impairment at regional and global levels for 2010 with trends from 1990. Pediatr Res 2013; 74 Suppl 1: 50–72. DOI: 10.1038/pr.2013.206.

3. Shankaran S, Laptook AR, Ehrenkranz RA, et al. Whole-body hypothermia for neonates with hypoxic-ischemic encephalopathy. N Engl J Med 2005; 353: 1574–1584. DOI: 10.1056/NEJMcps050929.

4. Thayyil S, Pant S, Montaldo P, et al. Hypothermia for moderate or severe neonatal encephalopathy in low-income and middle-income countries (HELIX): a randomised controlled trial in India, Sri Lanka, and Bangladesh. Lancet Glob Health 2021; 9: e1273–e1285. 20210803. DOI: 10.1016/S2214-109X(21)00264-3.

5. Hung P, Casey MM, Kozhimannil KB, et al. Rural-urban differences in access to hospital obstetric and neonatal care: how far is the closest one? J Perinatol 2018; 38: 645–652. 20180216. DOI: 10.1038/s41372-018-0063-5.

6. Pires A, Fortuna A, Alves G and Falcao A. Intranasal drug delivery: how, why and what for? J Pharm Pharm Sci 2009; 12: 288–311. DOI: 10.18433/j3nc79.

7. Snyers D, Tribolet S and Rigo V. Intranasal Analgosedation for Infants in the Neonatal Intensive Care Unit: A Systematic Review. Neonatology 2022; 119: 273–284. 20220301. DOI: 10.1159/000521949.

8. Ku LC, Simmons C, Smith PB, et al. Intranasal midazolam and fentanyl for procedural sedation and analgesia in infants in the neonatal intensive care unit. J Neonatal Perinatal Med 2019; 12: 143–148. DOI: 10.3233/NPM-17149.

9. Kaushal S, Placencia JL, Maffei SR and Chumpitazi CE. Intranasal Fentanyl Use in Neonates. Hosp Pharm 2020; 55: 126–129. 20190204. DOI: 10.1177/0018578719828335.

10. Wolfe TR and Braude DA. Intranasal medication delivery for children: a brief review and update. Pediatrics 2010; 126: 532–537. 20100809. DOI: 10.1542/peds.2010-0616.

11. Grassin-Delyle S, Buenestado A, Naline E, et al. Intranasal drug delivery: an efficient and non-invasive route for systemic administration: focus on opioids. Pharmacol Ther 2012; 134: 366–379. 2012/04/03. DOI: 10.1016/j.pharmthera.2012.03.003.

12. Oneal RM, Beil Jr RJ and Schlesinger J. Surgical anatomy of the nose. Otolaryngol Clin North Am 1999; 32: 145–181. DOI: 10.1016/s0030-6665(05)70119-5.

13. Merkus FW, Verhoef JC, Schipper NG and Marttin E. Nasal mucociliary clearance as a factor in nasal drug delivery. Adv Drug Deliv Rev 1998; 29: 13–38. DOI: 10.1016/s0169-409x(97)00059-8.

14. Illum L. Nasal drug delivery--possibilities, problems and solutions. J Control Release 2003; 87: 187–198. DOI: 10.1016/s0168-3659(02)00363-2.

15. Gosau M, Rink D, Driemel O and Draenert FG. Maxillary sinus anatomy: a cadaveric study with clinical implications. Anat Rec (Hoboken) 2009; 292: 352–354. DOI: 10.1002/ar.20859.

16. Charlton S, Jones NS, Davis SS and Illum L. Distribution and clearance of bioadhesive formulations from the olfactory region in man: effect of polymer type and nasal delivery device. Eur J Pharm Sci 2007; 30: 295–302. 20061210. DOI: 10.1016/j.ejps.2006.11.018.

17. Crowe TP, Greenlee MHW, Kanthasamy AG and Hsu WH. Mechanism of intranasal drug delivery directly to the brain. Life Sci 2018; 195: 44–52. 20171222. DOI: 10.1016/j.lfs.2017.12.025.

18. Vyas TK, Shahiwala A, Marathe S and Misra A. Intranasal drug delivery for brain targeting. Curr Drug Deliv 2005; 2: 165–175. DOI: 10.2174/1567201053586047.

19. Miyake MM, Nocera A and Miyake MM. P-glycoprotein and chronic rhinosinusitis. World J Otorhinolaryngol Head Neck Surg 2018; 4: 169-174. 20180824. DOI: 10.1016/j.wjorl.2018.07.002.

20. Pardridge WM. Blood-brain barrier biology and methodology. J Neurovirol 1999; 5: 556-569. DOI: 10.3109/13550289909021285.

21. Kristensson K and Olsson Y. Uptake of exogenous proteins in mouse olfactory cells. Acta Neuropathol 1971; 19: 145–154. DOI: 10.1007/BF00688493.

22. Jansson B and Bjork E. Visualization of in vivo olfactory uptake and transfer using fluorescein dextran. J Drug Target 2002; 10: 379–386. DOI: 10.1080/1061186021000001823.

23. Thorne RG and Frey WH, 2nd. Delivery of neurotrophic factors to the central nervous system: pharmacokinetic considerations. Clin Pharmacokinet 2001; 40: 907–946. DOI: 10.2165/00003088-200140120-00003.

24. Li Y, Field PM and Raisman G. Olfactory ensheathing cells and olfactory nerve fibroblasts maintain continuous open channels for regrowth of olfactory nerve fibres. Glia 2005; 52: 245–251. DOI: 10.1002/glia.20241.

25. Takano K, Kojima T, Go M, et al. HLA-DR- and CD11c-positive dendritic cells penetrate beyond well-developed epithelial tight junctions in human nasal mucosa of allergic rhinitis. J Histochem Cytochem 2005; 53: 611–619. DOI: 10.1369/jhc.4A6539.2005.

26. Kida S, Pantazis A and Weller RO. CSF drains directly from the subarachnoid space into nasal lymphatics in the rat. Anatomy, histology and immunological significance. Neuropathol Appl Neurobiol 1993; 19: 480–488. DOI: 10.1111/j.1365-2990.1993.tb00476.x.

27. Johnston M, Zakharov A, Papaiconomou C, et al. Evidence of connections between cerebrospinal fluid and nasal lymphatic vessels in humans, non-human primates and other mammalian species. Cerebrospinal Fluid Res 2004; 1: 2. 20041210. DOI: 10.1186/1743-8454-1-2.

28. Lipchock SV, Reed DR and Mennella JA. The gustatory and olfactory systems during infancy: implications for development of feeding behaviors in the high-risk neonate. Clin Perinatol 2011; 38: 627–641. 20111013. DOI: 10.1016/j.clp.2011.08.008.

29. Doherty TM, Hu A and Salik I. Physiology, Neonatal. StatPearls. Treasure Island (FL), 2023.

30. te Pas AB, Wong C, Kamlin CO, et al. Breathing patterns in preterm and term infants immediately after birth. Pediatr Res 2009; 65: 352–356. DOI: 10.1203/PDR.0b013e318193f117.

31. Doty RL, Deems DA, Frye RE, et al. Olfactory sensitivity, nasal resistance, and autonomic function in patients with multiple chemical sensitivities. Arch Otolaryngol Head Neck Surg 1988; 114: 1422–1427. DOI: 10.1001/archotol.1988.01860240072027.

32. Kahn A, Groswasser J, Sottiaux M, et al. Mechanisms of obstructive sleep apneas in infants. Biol Neonate 1994; 65: 235–239. DOI: 10.1159/000244058.

33. Moreira PI, Smith MA, Zhu X, et al. Oxidative stress and neurodegeneration. Ann N Y Acad Sci 2005; 1043: 545-552. DOI: 10.1196/annals.1333.062.

34. Gizurarson S. Anatomical and histological factors affecting intranasal drug and vaccine delivery. Curr Drug Deliv 2012; 9: 566–582. DOI: 10.2174/156720112803529828.

35. Chamanza R and Wright JA. A Review of the Comparative Anatomy, Histology, Physiology and Pathology of the Nasal Cavity of Rats, Mice, Dogs and Non-human Primates. Relevance to Inhalation Toxicology and Human Health Risk Assessment. J Comp Pathol 2015; 153: 287–314. 20151014. DOI: 10.1016/j.jcpa.2015.08.009.

36. Negus VE. [Observations on the comparative anatomy and physiology of olfaction]. Acta Otolaryngol 1954; 44: 13–24. DOI: 10.3109/00016485409126874.

37. Baak LM, Wagenaar N, van der Aa NE, et al. Feasibility and safety of intranasally administered mesenchymal stromal cells after perinatal arterial ischaemic stroke in the Netherlands (PASSIoN): a first-in-human, open-label intervention study. Lancet Neurol 2022; 21: 528–536. 2022/05/15. DOI: 10.1016/S1474-4422(22)00117-X.

38. Jalali R, Nogueira-Rodrigues A, Das A, et al. Drug Development in Low- and Middle-Income Countries: Opportunity or Exploitation? Am Soc Clin Oncol Educ Book 2022; 42: 1–8. DOI: 10.1200/EDBK_10033.

39. McDermott C CN. Prehospital Medication Administration: A Randomised Study Comparing Intranasal and Intravenous Routes. Emerg Med Int 2012; 2012: 5.

40. Inokuchi R O-FN, Nakamura K, Wada T, Gunshin M, Kitsuta Y, Nakajima S, Yahagi N. Comparison of Intranasal and Intravenous Diazepam on Status Epilepticus in Stroke Patients. Medicine 2015; 94.

41. Ben Abdelaziz R, Hafsi H, Hajji H, et al. Peripheral venous catheter complications in children: predisposing factors in a multicenter prospective cohort study. BMC Pediatr 2017; 17: 208. 20171219. DOI: 10.1186/s12887-017-0965-y.

42. Lemyre B and Chau V. Hypothermia for newborns with hypoxic-ischemic encephalopathy. Paediatr Child Health 2018; 23: 285–291. 20180612. DOI: 10.1093/pch/pxy028.

43. Suryanto, Plummer V and Boyle M. EMS Systems in Lower-Middle Income Countries: A Literature Review. Prehosp Disaster Med 2017; 32: 64–70. 20161212. DOI: 10.1017/S1049023X1600114X.

44. Robertson NJ, Meehan C, Martinello KA, et al. Human umbilical cord mesenchymal stromal cells as an adjunct therapy with therapeutic hypothermia in a piglet model of perinatal asphyxia. Cytotherapy 2021; 23: 521–535. 20201128. DOI: 10.1016/j.jcyt.2020.10.005.

45. van Velthoven CT, Kavelaars A, van Bel F and Heijnen CJ. Nasal administration of stem cells: a promising novel route to treat neonatal ischemic brain damage. Pediatr Res 2010; 68: 419–422. DOI: 10.1203/PDR.0b013e3181f1c289.

46. Donega V, Nijboer CH, Braccioli L, et al. Intranasal administration of human MSC for ischemic brain injury in the mouse: in vitro and in vivo neuroregenerative functions. PLoS One 2014; 9: e112339. 20141114. DOI: 10.1371/journal.pone.0112339.

47. Donega V, Nijboer CH, van Tilborg G, et al. Intranasally administered mesenchymal stem cells promote a regenerative niche for repair of neonatal ischemic brain injury. Exp Neurol 2014; 261: 53–64. 20140616. DOI: 10.1016/j.expneurol.2014.06.009.

48. Donega V, van Velthoven CT, Nijboer CH, et al. Intranasal mesenchymal stem cell treatment for neonatal brain damage: long-term cognitive and sensorimotor improvement. PLoS One 2013; 8: e51253. 20130103. DOI: 10.1371/journal.pone.0051253.

49. Jain A, Kratimenos P, Koutroulis I, et al. Effect of Intranasally Delivered rh-VEGF165 on Angiogenesis Following Cerebral Hypoxia-Ischemia in the Cerebral Cortex of Newborn Piglets. Int J Mol Sci 2017; 18 20171107. DOI: 10.3390/ijms18112356.

50. Lin S, Fan LW, Rhodes PG and Cai Z. Intranasal administration of IGF-1 attenuates hypoxic-ischemic brain injury in neonatal rats. Exp Neurol 2009; 217: 361–370. 20090328. DOI: 10.1016/j.expneurol.2009.03.021.

51. Scafidi J, Hammond TR, Scafidi S, et al. Intranasal epidermal growth factor treatment rescues neonatal brain injury. Nature 2014; 506: 230–234. 20131225. DOI: 10.1038/nature12880.

52. Chung S, Peters JM, Detyniecki K, et al. The nose has it: Opportunities and challenges for intranasal drug administration for neurologic conditions including seizure clusters. Epilepsy Behav Rep 2023; 21: 100581. 20221228. DOI: 10.1016/j.ebr.2022.100581.

53. Linakis MW, Roberts JK, Lala AC, et al. Challenges Associated with Route of Administration in Neonatal Drug Delivery. Clin Pharmacokinet 2016; 55: 185–196. DOI: 10.1007/s40262-015-0313-z.

54. Martins D, Brodmann K, Veronese M, et al. "Less is more": A dose-response account of intranasal oxytocin pharmacodynamics in the human brain. Prog Neurobiol 2022; 211: 102239. 20220203. DOI: 10.1016/j.pneurobio.2022.102239.

55. Pietrowsky R, Struben C, Molle M, et al. Brain potential changes after intranasal vs. intravenous administration of vasopressin: evidence for a direct nose-brain pathway for peptide effects in humans. Biol Psychiatry 1996; 39: 332–340. DOI: 10.1016/0006-3223(95)00180-8.

56. Smith PF. Neuroprotection against hypoxia-ischemia by insulin-like growth factor-I (IGF-I). IDrugs 2003; 6: 1173–1177.

57. Walters EM, Wells KD, Bryda EC, et al. Swine models, genomic tools and services to enhance our understanding of human health and diseases. Lab Anim (NY) 2017; 46: 167–172. DOI: 10.1038/laban.1215.

58. Koehler RC, Yang ZJ, Lee JK and Martin LJ. Perinatal hypoxic-ischemic brain injury in large animal models: Relevance to human neonatal encephalopathy. J Cereb Blood Flow Metab 2018; 38: 2092–2111. 20180828. DOI: 10.1177/0271678X18797328.

59. Gieling ET ST, Nordquist R, van der Staay FJ. The pig as a model animal for studying cognition and neurobehavioral disorders. Curr Top Behav Neurosci 2011; 7: 83.

60. Chavez-Valdez RB, J.; Carlin, K. Chapter 12 - Rodent modeling of neonatal hypoxic-ischemic brain injury. Handbook of Animal Models in Neurological Disorders 2023.

61. Uslu ZSA. Recent advancements in behavioral testing in rodents. MethodsX 2021; 8: 101536. 20210928. DOI: 10.1016/j.mex.2021.101536.

62. Burnsed JC, Chavez-Valdez R, Hossain MS, et al. Hypoxia-ischemia and therapeutic hypothermia in the neonatal mouse brain--a longitudinal study. PLoS One 2015; 10: e0118889. 20150316. DOI: 10.1371/journal.pone.0118889.

63. Diaz J, Abiola S, Kim N, et al. Therapeutic Hypothermia Provides Variable Protection against Behavioral Deficits after Neonatal Hypoxia-Ischemia: A Potential Role for Brain-Derived Neurotrophic Factor. Dev Neurosci 2017; 39: 257–272. 20170215. DOI: 10.1159/000454949.

64. Patel SD, Pierce L, Ciardiello A, et al. Therapeutic hypothermia and hypoxia-ischemia in the term-equivalent neonatal rat: characterization of a translational preclinical model. Pediatr Res 2015; 78: 264–271. 20150521. DOI: 10.1038/pr.2015.100.

65. Vannucci RC and Vannucci SJ. Perinatal hypoxic-ischemic brain damage: evolution of an animal model. Dev Neurosci 2005; 27: 81–86. DOI: 10.1159/000085978.

66. McDonald CA, Djuliannisaa Z, Petraki M, et al. Intranasal Delivery of Mesenchymal Stromal Cells Protects against Neonatal Hypoxic(-)Ischemic Brain Injury. Int J Mol Sci 2019; 20 20190517. DOI: 10.3390/ijms20102449.

67. Ji G, Liu M, Zhao XF, et al. NF-kappaB Signaling is Involved in the Effects of Intranasally Engrafted Human Neural Stem Cells on Neurofunctional Improvements in Neonatal Rat Hypoxic-Ischemic Encephalopathy. CNS Neurosci Ther 2015; 21: 926–935. 20150808. DOI: 10.1111/cns.12441.

68. Lu S, Li K, Yang Y, et al. Optimization of an Intranasal Route for the Delivery of Human Neural Stem Cells to Treat a Neonatal Hypoxic-Ischemic Brain Injury Rat Model. Neuropsychiatr Dis Treat 2022; 18: 413–426. 20220223. DOI: 10.2147/NDT.S350586.

69. Lawson A, Snyder W and Peeples ES. Intranasal Administration of Extracellular Vesicles Mitigates Apoptosis in a Mouse Model of Neonatal Hypoxic-Ischemic Brain Injury. Neonatology 2022; 119: 345–353. 20220325. DOI: 10.1159/000522644.

70. Turovsky EA, Golovicheva VV, Varlamova EG, et al. Mesenchymal stromal cell-derived extracellular vesicles afford neuroprotection by modulating PI3K/AKT pathway and calcium oscillations. Int J Biol Sci 2022; 18: 5345–5368. 20220821. DOI: 10.7150/ijbs.73747.

71. Moran J, Stokowska A, Walker FR, et al. Intranasal C3a treatment ameliorates cognitive impairment in a mouse model of neonatal hypoxic-ischemic brain injury. Exp Neurol 2017; 290: 74–84. 20170104. DOI: 10.1016/j.expneurol.2017.01.001.

72. Dixon BJ, Chen D, Zhang Y, et al. Intranasal Administration of Interferon Beta Attenuates Neuronal Apoptosis via the JAK1/STAT3/BCL-2 Pathway in a Rat Model of Neonatal Hypoxic-Ischemic Encephalopathy. ASN Neuro 2016; 8 20160928. DOI: 10.1177/1759091416670492.

73. Harding B, Conception K, Li Y and Zhang L. Glucocorticoids Protect Neonatal Rat Brain in Model of Hypoxic-Ischemic Encephalopathy (HIE). Int J Mol Sci 2016; 18 20161222. DOI: 10.3390/ijms18010017.

74. Nijboer CH, Heijnen CJ, Groenendaal F, et al. Strong neuroprotection by inhibition of NF-kappaB after neonatal hypoxia-ischemia involves apoptotic mechanisms but is independent of cytokines. Stroke 2008; 39: 2129–2137. 20080417. DOI: 10.1161/STROKEAHA.107.504175.

75. Yang D, Sun YY, Lin X, et al. Intranasal delivery of cell-penetrating anti-NF-kappaB peptides (Tat-NBD) alleviates infection-sensitized hypoxic-ischemic brain injury. Exp Neurol 2013; 247: 447–455. 20130123. DOI: 10.1016/j.expneurol.2013.01.015.

76. Yang D, Sun YY, Lin X, et al. Taming neonatal hypoxic-ischemic brain injury by intranasal delivery of plasminogen activator inhibitor-1. Stroke 2013; 44: 2623–2627. 20130723. DOI: 10.1161/STROKEAHA.113.001233.

77. Chen D, Dixon BJ, Doycheva DM, et al. IRE1alpha inhibition decreased TXNIP/NLRP3 inflammasome activation through miR-17-5p after neonatal hypoxic-ischemic brain injury in rats. J Neuroinflammation 2018; 15: 32. 20180202. DOI: 10.1186/s12974-018-1077-9.

78. Wang Z, Zhao H, Peng S and Zuo Z. Intranasal pyrrolidine dithiocarbamate decreases brain inflammatory mediators and provides neuroprotection after brain hypoxia-ischemia in neonatal rats. Exp Neurol 2013; 249: 74–82. 20130829. DOI: 10.1016/j.expneurol.2013.08.006.

79. Kuan CY, Chen HR, Gao N, et al. Brain-targeted hypoxia-inducible factor stabilization reduces neonatal hypoxic-ischemic brain injury. Neurobiol Dis 2021; 148: 105200. 20201126. DOI: 10.1016/j.nbd.2020.105200.

80. Shi X, Xu L, Doycheva DM, et al. Sestrin2, as a negative feedback regulator of mTOR, provides neuroprotection by activation AMPK phosphorylation in neonatal hypoxic-ischemic encephalopathy in rat pups. J Cereb Blood Flow Metab 2017; 37: 1447–1460. 20160101. DOI: 10.1177/0271678X16656201.

81. Xie S, Jiang X, Doycheva DM, et al. Activation of GPR39 with TC-G 1008 attenuates neuroinflammation via SIRT1/PGC-1alpha/Nrf2 pathway post-neonatal hypoxic-ischemic injury in rats. J Neuroinflammation 2021; 18: 226. 20211013. DOI: 10.1186/s12974-021-02289-7.

82. Yu S, Doycheva DM, Gamdzyk M, et al. Activation of MC1R with BMS-470539 attenuates neuroinflammation via cAMP/PKA/Nurr1 pathway after neonatal hypoxic-ischemic brain injury in rats. J Neuroinflammation 2021; 18: 26. 20210119. DOI: 10.1186/s12974-021-02078-2.

83. Ezenwa BN, Olorunfemi G, Fajolu I, et al. Trends and predictors of in-hospital mortality among babies with hypoxic ischaemic encephalopathy at a tertiary hospital in Nigeria: A retrospective cohort study. PLoS One 2021; 16: e0250633. 20210426. DOI: 10.1371/journal.pone.0250633.

84. Ross TM, Martinez PM, Renner JC, et al. Intranasal administration of interferon beta bypasses the blood-brain barrier to target the central nervous system and cervical lymph nodes: a non-invasive treatment strategy for multiple sclerosis. J Neuroimmunol 2004; 151: 66–77. DOI: 10.1016/j.jneuroim.2004.02.011.

85. Barr RD. The importance of lowering the costs of stem cell transplantation in developing countries. Int J Hematol 2002; 76 Suppl 1: 365–367. DOI: 10.1007/BF03165286.

86. Chakkarapani AA, Aly H, Benders M, et al. Therapies for neonatal encephalopathy: Targeting the latent, secondary and tertiary phases of evolving brain injury. Semin Fetal Neonatal Med 2021; 26: 101256. 20210612. DOI: 10.1016/j.siny.2021.101256.

87. Nijboer CH, Heijnen CJ, Groenendaal F, et al. A dual role of the NF-kappaB pathway in neonatal hypoxic-ischemic brain damage. Stroke 2008; 39: 2578–2586. 20080417. DOI: 10.1161/STROKEAHA.108.516401.

88. Deng C, Li J, Li L, et al. Effects of hypoxia ischemia on caspase-3 expression and neuronal apoptosis in the brain of neonatal mice. Exp Ther Med 2019; 17: 4517–4521. 20190415. DOI: 10.3892/etm.2019.7487.

89. Northington FJ, Ferriero DM, Graham EM, et al. Early Neurodegeneration after Hypoxia-Ischemia in Neonatal Rat Is Necrosis while Delayed Neuronal Death Is Apoptosis. Neurobiol Dis 2001; 8: 207–219. DOI: 10.1006/nbdi.2000.0371.

90. Northington FJ, Chavez-Valdez R, Graham EM, et al. Necrostatin decreases oxidative damage, inflammation, and injury after neonatal HI. J Cereb Blood Flow Metab 2011; 31: 178–189. 20100623. DOI: 10.1038/jcbfm.2010.72.

91. Wang X HW, Du X, Changlian Z, Carlsson Y, Mallard C, Jacotot E, Hagberg H. Neuroprotective effect of Bax-inhibiting peptide on neonatal brain injury. Stroke 2010; 41: 2050.

92. Levison SW R-FE, Kim BH, Hagberg H, Fleiss B, Gressens P, Dobrowolski R. Mechanisms of Tertiary Neurodegeneration after Neonatal Hypoxic-Ischemic Brain Damage. Pediatr Med 2022; 5: 20.

93. Nakajima W, Ishida A, Lange MS, et al. Apoptosis has a prolonged role in the neurodegeneration after hypoxic ischemia in the newborn rat. J Neurosci 2000; 20: 7994–8004. DOI: 10.1523/JNEUROSCI.20-21-07994.2000.

94. Wu YW, Comstock BA, Gonzalez FF, et al. Trial of Erythropoietin for Hypoxic-Ischemic Encephalopathy in Newborns. N Engl J Med 2022; 387: 148–159. DOI: 10.1056/NEJMoa2119660.

95. Nair J and Kumar VHS. Current and Emerging Therapies in the Management of Hypoxic Ischemic Encephalopathy in Neonates. Children (Basel) 2018; 5 20180719. DOI: 10.3390/children5070099.

96. The-World-Bank. GDP (current US$). 2022.

97. Reznik GK. Comparative anatomy, physiology, and function of the upper respiratory tract. Environ Health Perspect 1990; 85: 171–176. DOI: 10.1289/ehp.85-1568330.

98. Zhang YT, He KJ, Zhang JB, et al. Advances in intranasal application of stem cells in the treatment of central nervous system diseases. Stem Cell Res Ther 2021; 12: 210. 20210324. DOI: 10.1186/s13287-021-02274-0.

